# Outside-in engineering of cadherin endocytosis using a conformation strengthening antibody

**DOI:** 10.1101/2024.08.15.608168

**Authors:** Bin Xie, Shipeng Xu, Sanjeevi Sivasankar

## Abstract

P-cadherin, a crucial cell-cell adhesion protein which is overexpressed in numerous malignant cancers, is a popular target for drug delivery antibodies. However, molecular guidelines for engineering antibodies that can be internalized upon binding to P-cadherin are unknown. Here, we use a combination of biophysical, biochemical, and cell biological methods to demonstrate that trapping the cadherin extracellular region in an X-dimer adhesive conformation, triggers cadherin endocytosis via a novel outside-in signaling mechanism. We show that the monoclonal antibody CQY684 traps P-cadherin in an X-dimer conformation and strengthens this adhesive structure. Formation of stable X-dimers results in the dissociation of p120-catenin, a suppressor of cadherin endocytosis, from the X-dimer cytoplasmic region. This increases the turnover of P-cadherin and targets the cadherin-antibody complex to the lysosome. Our results establish a previously unknown outside-in signaling mechanism that provides fundamental insights into how cells regulate adhesion and that can be exploited by anti-cadherin antibodies for intracellular drug delivery.

## Introduction

P-cadherin (placental cadherin; Pcad), a member of the classical cadherin family of cell-cell adhesion proteins, is overexpressed in malignant cancers^1^, including cancers that originate from the breast^2^, pancreas^3^ and lung^4^. Consequently, Pcad is recognized as a promising monoclonal antibody (Mab) target for anti-cancer drug delivery. In therapeutic applications, drug conjugated Mabs specifically target the extracellular regions of adherent Pcads at cell-cell junctions. Upon binding to Pcad ectodomains, the Mabs are internalized for payload release. However, the molecular linkage between extracellular cadherin conformations and cadherin endocytosis is completely unknown. Consequently, molecular guidelines for engineering Mabs that bind to Pcad ectodomains and promote cadherin internalization have not been established.

Like all classical cadherins, Pcad ectodomains bind in two distinct adhesive conformations: X-dimers and strand-swap dimers (S-dimers)^5,6^. Due to their faster on-rate and weaker binding strength, X-dimers are thought to be an intermediate in the formation and dissociation of S- dimers^6–8^. Notably, cells expressing mutant cadherins trapped in an X-dimer conformation, exhibit more dynamic cell junction due to increased cadherin turnover^7^. However, the mechanism by which X-dimer formation promotes cadherin internalization is unknown.

A master regulator of cadherin internalization is p120-catenin (p120), a protein that binds to the cadherin cytoplasmic region^9^. By acting as an endocytosis inhibitor, p120 stabilizes cadherins on the cell surface. Conversely, the dissociation of p120 triggers cadherin endocytosis and promotes their internalization^10^. However, it is unclear if X-dimer formation triggers cytoplasmic p120 dissociation and promotes cadherin endocytosis. Studying this outside-in mechanism is challenging due to a lack of methods to trap cadherins in an X-dimer conformation without introducing mutations.

One strategy to trap cadherins in distinct binding conformations is the use of Mabs. For instance, two E-cadherin (Ecad) specific mAbs 19A11 and 66E8, have been shown to bind stabilize S-dimers^11–13^. The stabilization of Ecad S-dimers on the cell surface has been shown to strengthen cell adhesion and promote the dephosphorylation of p120^14,15^, suggesting a possible outside-in relationship between Ecad extracellular binding conformations and intracellular signaling. Conversely, the Mab CQY684, part of the antibody-drug conjugate PCA062, was recently shown to recognize the ectodomains of Pcad. This resulted in the endocytosis of the Mab-Pcad complex, thereby facilitating drug delivery^16^. Despite the potential of CQY684 as a drug delivery platform and its usage in Phase-I clinical trials^17^, the molecular mechanism by which CQY684 triggers Pcad endocytosis is unknown.

In this study, we combine biophysical (atomic force microscopy, and bead aggregation), computational (molecular dynamics and steered molecular dynamics simulations), biochemical (surface biotinylation, and co-immunoprecipitation), and cell biological (cell adhesion assays, confocal imaging, and fluorescence recovery after photobleaching) measurements to demonstrate that CQY684 traps Pcad in an X-dimer conformation and stabilizes this binding structure. We show that the stabilization of X-dimers triggers the dissociation of p120 from the Pcad cytoplasmic region which targets the antibody-Pcad complex to the lysosome. Our results establish a previously unknown outside-in signaling mechanism that provides fundamental insights into how cells regulate adhesion. Cadherin outside-in signaling can also be exploited by antibodies for intracellular drug delivery.

## Results

### CQY684 selectively strengthens Pcad X-dimers

Since a previous structural study has shown that CQY684 binding does not interfere with formation of either Pcad S-dimers or X-dimers^16^, we used atomic force microscopy (AFM) to measure the effect of CQY684 on the stability of these distinct Pcad conformations. We used three constructs in our experiments: human Pcad W2A mutant (trapped in an X-dimer conformation)^6^, human Pcad K14E mutant (trapped in an S-dimer conformation)^6^, and wild-type human Pcad (WT). We immobilized the complete extracellular region of each Pcad construct (EC1–5) on an AFM cantilever and glass substrate functionalized with polyethylene glycol (PEG) tethers, and measured Pcad–Pcad interactions with or without 40nM CQY684 in the buffer (‘+CQY’ or ‘−CQY’, Figure 1a, upper panel).

**Figure 1.**
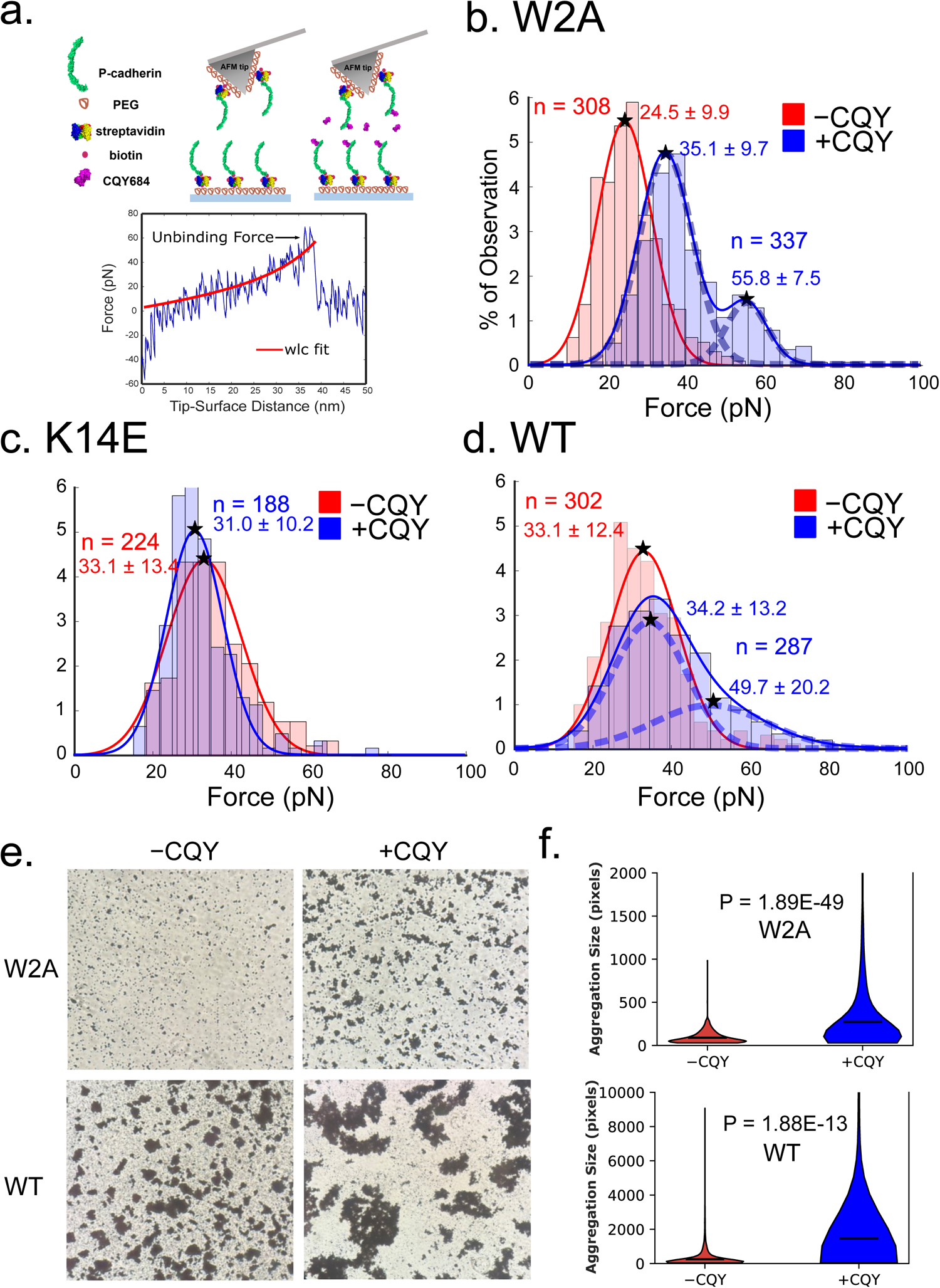
CQY684 Fab strengthens Pcad X-dimers. (a) Upper panel: scheme for AFM experiments carried out in the absence of CQY684 Fab (−CQY), and in the presence of CQY684 Fab (+CQY). Full ectodomains of Pcad were immobilized on AFM tips and substrates functionalized with PEG tethers. Bottom panel: example force curve. Stretching of the PEG tether, which serves as a “signature” of a single molecule unbinding event, was fit to a WLC model (red line). (b)(c)(d) AFM Experiments were performed with W2A, K14E and WT Pcad in the absence (−CQY, red curves and histograms) and presence (+CQY, blue curves and histograms) of CQY684 Fab. Histograms of the unbinding forces were generated by binning the data in each condition using the Freedman–Diaconis rule. The optimal number of Gaussian distributions for each fit was determined using BIC. (b) In -CQY condition, W2A Pcad unbinding forces were best described by a single Gaussian (peak at 24.5 ± 9.9 pN). Addition of CQY684 strengthened X-dimers; unbinding forces increased significantly and the distribution was best described by two Gaussians at a higher peak force (35.1 ± 9.7 pN and 55.8 ± 7.5 pN respectively). (c) Unbinding force distributions measured with K14E Pcad were similar in both the -CQY and +CQY condition demonstrating that CQY684 did not strengthen S-dimers. (d) Unbinding force distributions measured with WT Pcad in the absence of CQY684 was best fit by a single Gaussian (peak force of 33.1 ± 12.4 pN corresponding to S-dimer conformation). Addition of CQY684 selectively strengthened the X-dimer and resulted in a bimodal force distribution (peaks at 34.2 ± 13.2 pN and 49.7 ± 20.2 pN respectively). (e) Bead aggregation assays showed that beads functionalized with either W2A or WT Pcad formed larger aggregates in the presence of CQY684 Fab. (f) Volin plots of bead aggregate sizes observed in different conditions. Mean size of the aggregate is show as a black line on each violin. Two independent biological repeats were performed for bead aggregation experiments. Student t-test was performed on the violin plot.

During a typical AFM measurement, the cantilever and substrate, both functionalized with Pcad, were brought into contact to allow cadherin interactions. The tip was then retracted at a constant velocity (1 µm/s) to measure the force required to rupture the Pcad–Pcad bond. Unbinding events, characterized by nonlinear stretching of PEG tethers, were analyzed using a worm-like chain (WLC) model with least-squares fitting (Figure 1a, lower panel). We confirmed CQY684’s recognition of all mutants using Western blots (Supplementary Figure S1). Unbinding force histograms were fitted to Gaussian distributions, with the optimal number of distributions predicted using the Bayesian Information Criterion (Supplementary Figure S2).

For Pcad W2A mutants which can only form X-dimers, a single Gaussian distribution with a peak force of 24.5 ± 9.9 pN was observed in the −CQY condition. In the +CQY condition, W2A binding was strengthened and two Gaussian distributions with peak forces of 35 ± 9.7 pN and 55.7 ± 7.5 pN were observed (Figure 1b, Supplementary Figure S2a), indicating that CQY684 enhances Pcad X-dimer binding strength. Steered Molecular Dynamics simulations (described in the next section) provide an explanation for the bimodal force distribution measured in the W2A +CQY condition. The simulations suggest that besides stabilizing the X-dimer interface, CQY684 binding also stochastically forms a salt bridge between the two partner cadherins, which further strengthens X-dimer adhesion. Since this salt-bridge formation is probabilistic, it results in a bimodal distribution of unbinding forces.

For Pcad K14E mutants, which can only form S-dimers, single Gaussian distributions were observed in both −CQY and +CQY conditions with similar peak forces (33.1 ± 13.4 pN and 32.0 ± 10.2 pN, respectively (Figure 1c, Supplementary Figure S2b), suggesting that CQY684 does not affect Pcad S-dimer binding strength. Although WT cadherin ectodomains can adopt either conformation in solution, X-dimers are believed to serve as a transient intermediate and cadherins are known to ultimately adopt a stable S-dimer structure. Indeed, for Pcad WT in the -CQY condition, we observed a single Gaussian force distribution with a peak force of 33.0 ± 12.4 pN (Figure 1d , Supplementary Figure S2c upper panel), similar to Pcad K14E interactions but higher than Pcad W2A interactions, suggesting that Pcad WT primarily forms S-dimers. However, in the presence of CQY684, two Gaussian force distributions were observed with peaks at 34.2 ± 13.2 pN and 49.7 ± 20.2 pN (Figure 1d, Supplementary Figure S2c lower panel), similar to the force distribution of Pcad W2A in the +CQY condition. Since CQY684 only strengthens X-dimers and does not affect S-dimers, we concluded that the observed strengthening in the WT +CQY condition (Figure 1d) was due to CQY684 trapping the WT Pcad in an X-dimer structure.

In addition to AFM experiments, we conducted bead aggregation assays on the three Pcad constructs, with and without CQY684. While AFM experiments provide precise measurements of individual interactions, bead aggregation assays assess how CQY684 affects Pcad ectodomain binding in a more collective manner. In both Pcad W2A mutants and WT, the addition of CQY684 increased the size of the aggregates (Figure 1e, f), suggesting that CQY684 enhances Pcad-adhesion when the X-dimer state is accessible. The size of bead aggregates in Pcad WT +CQY was significantly larger than Pcad W2A +CQY due to the formation of both WT S-dimers and stabilized WT X-dimers. Consistent with previous studies^5,6^, Pcad K14E mutants did not form bead aggregates, likely due to the extremely low on-rate of K14E mutants, and the addition of CQY684 did not improve K14E aggregation (Supplementary Figure S3).

In summary, our cell-free experiments revealed that CQY684 selectively strengthens Pcad X-dimers without affecting S-dimers. The observed strengthening effect with WT cadherin indicates that CQY684 traps Pcad in a X-dimer structure.

### Molecular mechanism of CQY684 mediated stabilization of X-dimers

The crystal structure of CQY684 bound to the Pcad EC1-2 domain demonstrates that Pcad bound to the antibody adopts an X-dimer conformation (Figure 2a, upper panel). This is not surprising since the Pcads in the crystal structure possess an N-terminal extension and preferentially form X-dimers^6^. CQY684 directly binds to three loops of the Pcad EC1 domain (Figure 2b), referred to as ‘loop1’ (residues 15-17), ‘loop2’ (residues 43-51), and ‘loop3’ (residues 63-65). The binding interface of CQY684 is situated between the Pcad X-dimer interface, which is composed of residues 5-14, 20-23, 99-106, 138-145, and 196-204. Our previous studies have shown that antibodies binding near cadherin binding interfaces can stabilize these interfaces, thereby strengthening adhesion^11,12^. To test if CQY684 stabilizes the X-dimer binding interface, we performed molecular dynamics (MD) simulations on Pcad X-dimers under two conditions: ‘+CQY’ with two CQY684 Fabs bound to each Pcad in an X-dimer conformation (PDB 6ZTR), and ‘-CQY’ obtained from the same crystal structure by removing the CQY684 Fabs (Figure 2a).

**Figure 2.**
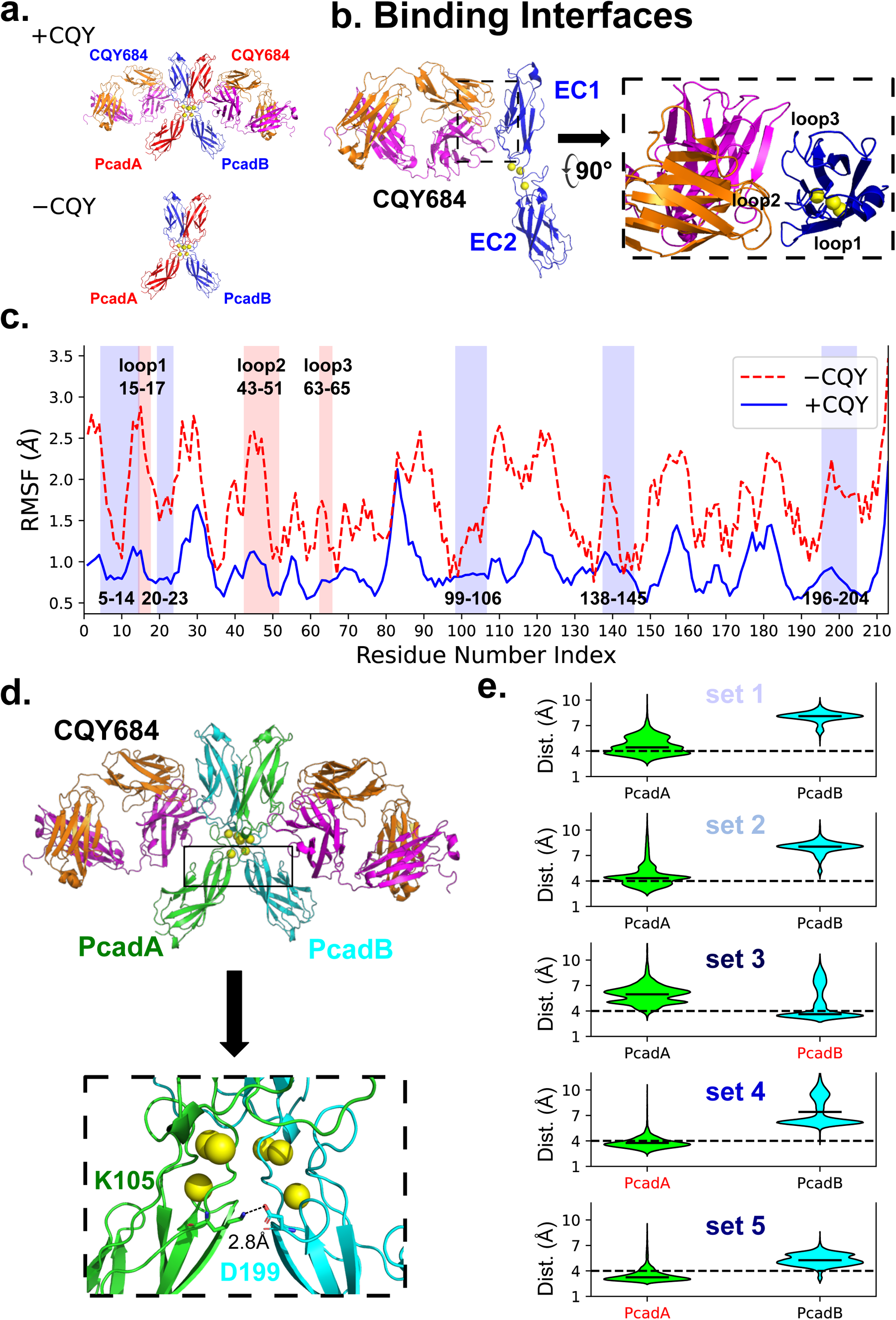
CQY684 stabilizes Pcad X-dimer binding interface. (a) Simulation setup for the +CQY conditions (with two CQY684 Fabs bound to an X-dimer) and the −CQY Pcad X-dimer. (b) CQY684 Fab binds to the Pcad EC1 domain by interacting with three loops. (c) Comparison of average RMSF values for Pcad residues 1-213 in the −CQY condition (red dashed line) and in the +CQY condition (blue solid line). The CQY684 binding interface is highlighted with a red background while the X-dimer interface is highlighted using a blue background. The lower RMSF values in the +CQY condition shows that the binding of CQY684 stabilizes the three loops of Pcad and also stabilizes the X-dimer binding interface. (d) CQY684 binding induces formation of a novel salt bridge between K105 and D199, within the X-dimer binding interface. The representative structure shown is the final structure of +CQY set 5. (e) Violin plots of distances between the charged atoms in the 105LYS:199ASP salt bridge measured during the last 40 ns of each +CQY MD simulations. The median distance is shown as a blue line on each violin. If the median distance is below the 4Å (black dashed line), then the salt bridge is considered to be stably formed during MD simulations. Salt bridge donor on PcadA is shown on the left while salt bridge donor on PcadB is shown on the right. Set 3 has one stable salt bridge with the donor from PcadB, while sets 4 and 5 each have one stable salt bridge with the donor from PcadA, all highlighted in red. No salt bridge formation was observed for sets 1 and 2.

Our MD simulations, set up as previously described^11,12^, included five independent repeats for each condition, with each repeat running for 60 ns until the system reached equilibrium (Supplementary Figure S4). To evaluate whether CQY684 stabilizes the Pcad X-dimer, we measured the root mean square fluctuations (RMSF) of the α-carbon residues during the last 30 ns of all MD simulations (Figure 2C). The average RMSF for Pcad with and without the antibody showed that CQY684 binding not only reduced the RMSF in its binding regions on Pcad (Figure 2C, red regions) but also stabilized the entire Pcad EC1 and EC2 domains, including the X-dimer binding interface (Figure 2C, blue regions).

Additionally, we characterized the electrostatic interactions (hydrogen bonds or salt bridges) between Pcads in all simulations (Supplementary Figure S5). Although CQY684 binding did not induce a gross conformational change in Pcad X-dimers, we observed that persistent interactions between Pcads (i.e. interactions that existed for at least 40% of simulation time), changed upon CQY684 binding. Significantly, a previously non-persistent salt bridge which was present in the X-dimer binding interface (105Lys:199Asp), became persistent (Figure 2d). To monitor the formation of this salt bridge, we calculated the distance between charged atoms. Using the criterion that salt bridges form when the median distance between charged atoms is below 4Å, our analysis showed that in the presence of CQY684 the 105Lys:199Asp salt bridge formed in three sets of simulations (sets 3-5, Figure 2e). Steered Molecular Dynamics (SMD) simulations (described below) showed that persistent formation of this salt bridge further strengthened Pcad X-dimers.

Although the conversion between cadherins X-dimers and S-dimers is not fully understood, we have previously shown that external pulling forces can convert X-dimers into S-dimers^18^. To test if the stabilized X-dimer induced by CQY684 binding inhibits this conversion, we performed SMD simulations on all the equilibrium structures from the MD simulations. In our SMD setup, we fixed the position of one Pcad and applied a constant force to pull the C-terminal of the other Pcad until the structures fully dissociated. To monitor the conversion of an X-dimer to an S-dimer structure, we calculated the root mean square deviation (RMSD) of the SMD structures relative to the Pcad S-dimer crystal structure (PDB code: 4ZMN). Independent of CQY684 binding, all X-dimers showed a conversion towards S-dimers before fully dissociating, upon application of pulling force (Figure 3a, b). However, in the absence of CQY684, the average conversion time was 1012 ± 212 ps, while CQY684 binding dramatically slowed this conversion nearly five-fold, with an average conversion time of 5307 ± 2355 ps. Given that previous studies suggest cadherin X-dimers are a necessary intermediate for S-dimer formation^5,6,19,20^, and our results show CQY684 inhibits the Pcad X-dimer transition to S-dimer, it is likely that CQY684 traps the interacting Pcad in an X-dimer conformation. This could explain why AFM measurements show that Pcad WT +CQY has a force distribution similar to W2A +CQY but different from K14E +CQY (Figure 1b-d).

**Figure 3.**
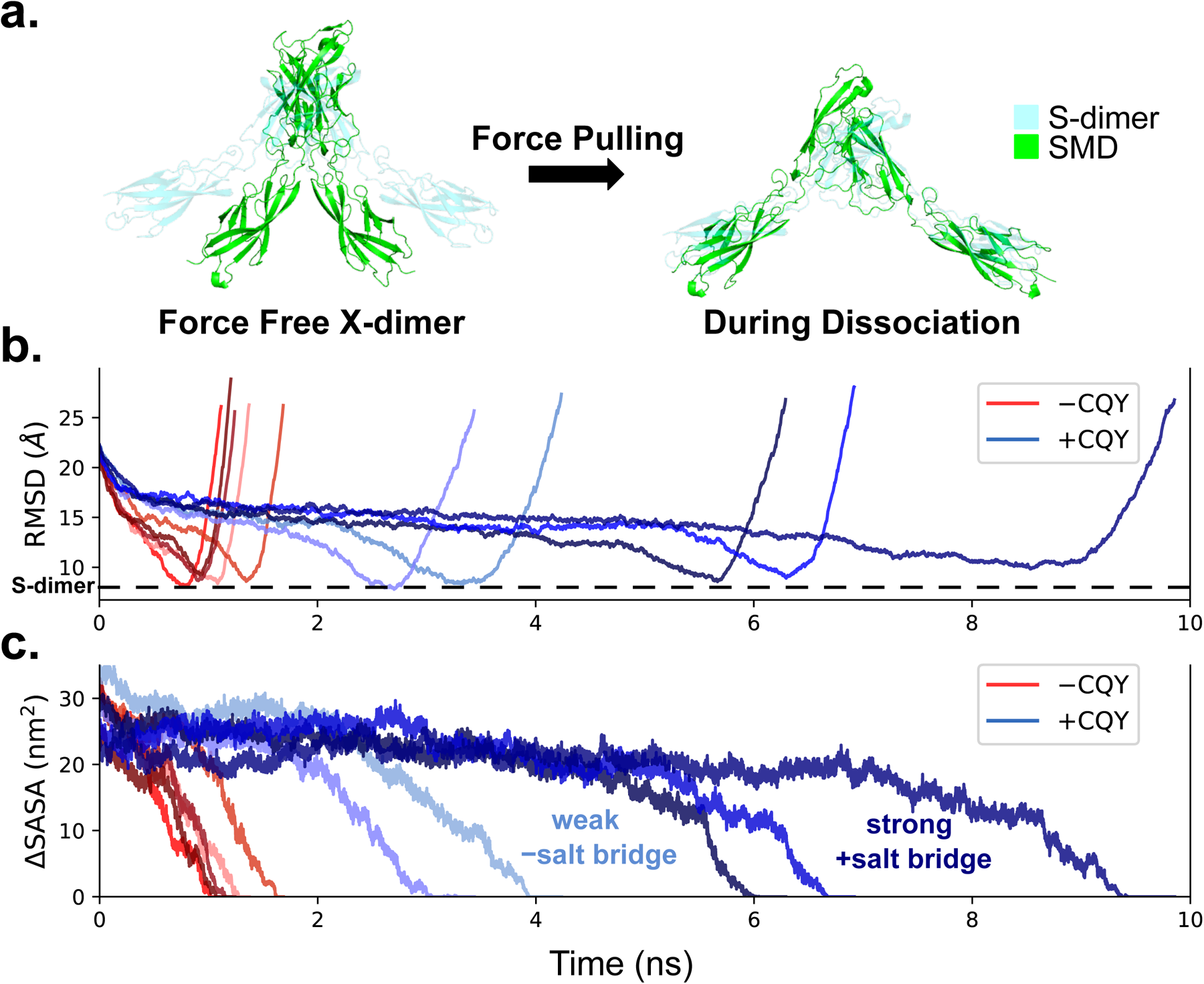
CQY684 inhibits Pcad transition from X-dimer to S-dimer and strengthens adhesion. (a) While dissociating under a constant pulling force, Pcad X-dimers convert to S-dimers. Selected structures during SMD are showed in green. The S-dimer reference crystal structure is showed in cyan. (b) RMSD was calculated between every SMD structure and the reference S-dimer crystal structure. The RMSD values first drop (indicating that the X-dimer converts to an S-dimer like structure) and then increase (as the dimer dissociates). The addition of CQY684 (different shades of blue) dramatically slows down this structural conversion compared to the −CQY condition (different shades of red). (c) Pcad unbinding during the constant force SMD simulation was calculated from changes in interfacial area computed from the ΔSASA, in the absence of CQY684 (shades of red), and in the presence of CQY684 (shades of blue). The addition of CQY684 stabilizes the Pcad X-dimers and slows down their dissociation. When the salt bridge forms during the MD simulation (in sets 3-5, highlighted in darker blue), the Pcad interactions are significantly stronger compared to conditions (sets 1 and 2, highlighted in lighter blue), where the salt bridge does not form. Color schemes for the +CQY conditions are the same as Figure 2e.

To test whether the stabilized X-dimer induced by CQY684 binding could lead to stronger interactions as observed in the AFM experiments, we measured the X-dimer dissociation time during each SMD simulation. Specifically, we estimated the interfacial binding area between the two Pcads during SMD by calculating the change in solvent accessible surface area (ΔSASA), where a decrease in ΔSASA to zero corresponds to the rupture of the interacting dimer (Figure 3c). In the absence of CQY684, the average dissociation time was 1207 ± 204 ps, while in the presence of CQY684, the average dissociation time increased to 5762 ± 2482 ps. Notably, when the novel salt bridge (105Lys:199Asp) stably formed between Pcads in sets 3-5, the average dissociation time of sets 3-5 (∼7300 ps, Figure 3c, dark blue) was almost two-fold longer than that of sets 1 and 2 (∼3500 ps, Figure 3c, light blue). This could explain why there are two gaussian distributions in the W2A + CQY and WT +CQY conditions for the AFM experiments (Figure 1b and d) with the lower force distribution corresponding to instances where the salt bridge is not formed and the higher force distribution corresponding to cases when the salt bridge is formed.

In summary, our computational studies revealed that CQY684 binding stabilizes the Pcad X-dimer conformation, preventing its conversion to an S-dimer structure. Furthermore, the stabilized X-dimer results in much stronger interactions, consistent with our AFM and bead aggregation results.

### CQY684 promotes Pcad endocytosis and disrupts Pcad-mediated cell adhesion

Cell-cell adhesion plays a crucial role in cancer progression; downregulation of cell adhesion occurs in cancer metastasis^21^ while upregulation of adhesion prevents metastasis^22^. However, the impact of CQY684 on Pcad-mediated cell-cell adhesion has not been previously characterized. Given that CQY684 strengthens Pcad X-dimers in solution and enhances bead aggregation with Pcad WT and W2A mutants, we anticipated that CQY684 would also enhance Pcad-mediated cell-cell adhesion. To test this, we performed cell adhesion assays on cells expressing either WT or W2A mutant Pcad. Specifically, we rescued Ecad and Pcad knockout (KO) A431 cells^23^ with either full length WT-Pcad or W2A-Pcad with an mCherry tag on the Pcad C-terminus.

To test the effect of CQY684 on Pcad-mediated cell-cell adhesion, we performed a dispase assay to evaluate cell adhesion strength under mechanical stress. In this assay, the cells expressing either WT-Pcad or W2A mutants were grown to 95% confluence, with 200 nM CQY684 Fab added during growth for the +CQY conditions. The cell monolayers were then detached from the dish using dispase enzyme and subjected to mechanical stress by vigorously rotating them on an orbital rotator for 2 hours. This process applied shear forces to the cell layers, allowing us to assess how adherent the cells remained under stress, both in the absence and presence of CQY684. Surprisingly, the monolayers treated with CQY684 showed significantly more fragmentation compared to those without CQY684 treatment (Figure 4a). Contrary to our expectations based on the bead aggregation results, the addition of CQY684 weakened cell-cell adhesion rather than strengthening it.

**Figure 4.**
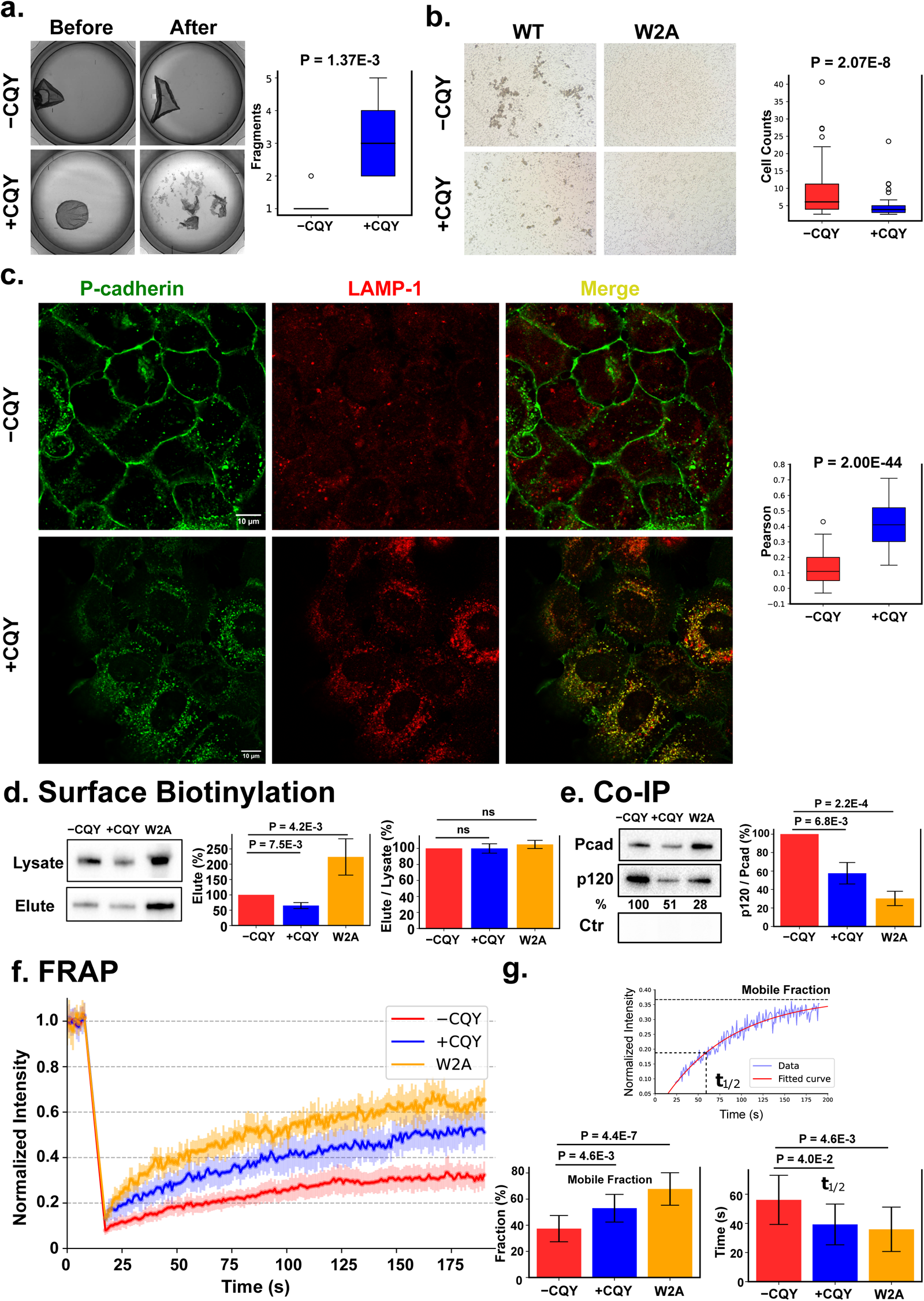
CQY684 disrupts Pcad mediated cell-cell adhesion by dissociating p120 catenin and promoting endocytosis. (a) Dispase assay shows that the addition of CQY684 Fab disrupts cell adhesion, and cell sheet monolayers become more fragmented. N = 10 for each condition, distributed across three independent biological repeats. (b) Cell aggregation assay shows that the addition of CQY684 Fab disrupts cell adhesion in the A431 cells expressing WT Pcad. A431 cells expressing W2A Pcad mutants do not form aggregates in both presence and absence of CQY684. Measurements performed across three biological repeats. (c) Immunofluorescence confocal imaging shows that, in the presence of CQY684, WT-Pcad colocalizes with the lysosomal marker LAMP1. Pcad is stained in green and LAMP1 is stained in red. Colocalization of Pcad and LAMP1 (yellow) is seen in the +CQY condition. N = 163 for ‘WT −CQY’ condition, and N = 131 for the ‘WT +CQY’ condition across three biological repeats. (d) Surface biotinylation assays. Left panel: Representative data. Total amount of Pcad in the cell lysate, and biotinylated surface Pcad was measured from the western blot signal intensity. Similar total protein amounts (as measured with a DC protein assay) was loaded in each lane. Middle panel: Bar-plot of the amount of cell surface Pcad (‘elute’) compared to the −CQY conditions. The amount of surface protein in the WT cells was reduced upon addition of CQY684 likely due to increased internalization of Pcad and subsequent lysosomal proteolysis. W2A cells exhibited higher levels of surface Pcad compared to WT cells due to higher transfection efficiency. Right panel: Bar-plot of surface Pcad versus the total amount of Pcad (elute / lysate) compared to the −CQY conditions. Differences between the three conditions are non-significant. Three independent repeats were performed in surface biotinylation assays. (e) Co-immunoprecipitation (Co-IP) experiments show that the amount of p120 association drops in X-dimers (i.e. either upon addition of CQY684 or in the W2A mutants). Left panel: Representative Co-IP data. The same amount of total protein (as measured with a DC protein assay) was loaded into each lane. Total amount of Pcad in the cells and the total amount of associated p120 were determined from the western blot signal intensity. No Pcad was detected when using an IgG control. Right panel: Bar-plot of three replicates. All values are compared to the −CQY conditions. (f) FRAP experiments were performed on cell-cell junctions in WT −CQY (red line), WT +CQY (blue line), and W2A (orange line) conditions. Normalized, average fluorescence signal of Pcad tagged with mCherry was measured. Both fluorescence recovery rate and mobile fraction increased in +CQY and W2A conditions, indicating more dynamic cell-cell junctions. Error bars for each data point show 95% confidence intervals. N = 20 for each condition measured across three biological repeats. (g) Upper panel: the fluorescence recovery was fitted to a single exponential model. Bottom left: comparison of the mobile fraction across the three conditions. Bottom right: comparison of the half recovery time across the three conditions. Student t-test was performed on all the box/bar plots.

In addition to the dispase assay, we also evaluated cell-cell adhesion using a cell aggregation assay. Consistent with the dispase assay results, A431 cells expressing WT-Pcad formed smaller aggregates in the presence of CQY684 (Figure 4b), indicating that CQY684 disrupts Pcad-mediated cell-cell adhesion. Notably, A431 cells expressing Pcad W2A mutants showed extremely low cell adhesion in both assays: in the dispase assay, cells could not be lifted as a single monolayer, and in the cell aggregation assay, no aggregates were observed with or without CQY684 (Figure 4b). The loss of adhesion in Pcad W2A mutants is likely due to the highly dynamic junctions formed by these mutants^7^.

To understand why CQY684 disrupts cell adhesion while strengthening the Pcad X-dimer conformation, we hypothesized that CQY684 traps WT-Pcad in an X-dimer conformation, which consequently enhances Pcad turnover. An indication of increased cadherin turnover is the internalization of cadherins and their subsequent colocalization with lysosomes. To test this hypothesis, we performed immunofluorescence imaging using a confocal microscope. A431 cells expressing WT-Pcad were grown to 70% confluency, with 200 nM CQY684 Fab added during cell growth for the +CQY condition. To assess Pcad colocalization with lysosomes, we stained for both Pcad and lysosomal-associated membrane protein 1 (Lamp1), a well-established lysosomal marker^24^. Cells treated with CQY684 Fab showed increased Pcad internalization and high colocalization with Lamp1 (Figure 4c), suggesting that CQY684 induces Pcad turnover, and lowers Pcad surface expression levels, thereby disrupting Pcad-mediated cell-cell adhesion.

To investigate changes in Pcad surface expression, we performed surface biotinylation experiments with WT -CQY, WT +CQY, and W2A cells. DC assays were used to normalize the protein amounts for the western blot analysis. As expected, the surface level of Pcad in WT cells decreased upon CQY684 treatment (Figure 4d, middle panel), likely due to increased internalization of Pcad and subsequent lysosomal proteolysis. While the W2A cells exhibited higher levels of surface Pcad compared to WT cells due to higher transfection efficiency (Figure 4d, middle panel), cell-cell adhesion between W2A cells was severely impaired. This suggests that factors beyond surface Pcad levels are responsible for adhesion defects in both WT +CQY and W2A conditions. Notably, the fraction of total Pcad expressed on the cell was similar (∼40% of total Pcad) across WT -CQY, WT +CQY, and W2A conditions (Figure 4d, right panel). Consistent with previous results^7^, this suggested that neither the addition of CQY684 nor the W2A mutation affected Pcad localization to the cell membrane.

### p120-catenin dissociates from the X-dimer cytoplasmic region, increasing junction dynamics

Previous studies have shown that p120 dissociation from the cadherin cytoplasmic domain signals endocytosis and increases cadherin turnover^10,25^. We therefore hypothesized that CQY684 binding would result in the dissociation of p120 from the Pcad cytoplasmic domain. In addition, given our previous findings that CQY684 traps Pcad in an X-dimer conformation, we hypothesized that CQY684-independent X-dimer formation would also trigger p120 dissociation from the cadherin cytoplasmic tail. To test this hypothesis, we conducted co-immunoprecipitation (Co-IP) experiments to measure the Pcad-p120 association in WT −CQY, WT +CQY, and W2A conditions (Figure 4e). In the Co-IP experiments, we used an anti-mCherry antibody to pull down Pcad as the bait and then measured the amount of p120 that co-precipitated with Pcad, normalizing these amounts to the WT −CQY condition. The WT +CQY showed ∼60% of p120 associated with Pcad compared to the WT -CQY condition, a 40% decrease in p120 binding (Figure 4e, right panel).. Similarly, only ∼30% of p120 were associated with the W2A mutant compared to the WT -CQY condition, confirming our hypothesis that X-dimer formation triggers p120 dissociation from the cadherin cytoplasmic tail (Figure 4e, right panel). The measured change in p120 association/dissociation did not arise from differential Pcad expression, because the surface biotinylation assays show that similar fraction of Pcad localize on the cell surface in all the conditions (Figure 4d, right panel).

To further understanding the effect of p120 dissociation and directly assess dynamics at cell-cell junctions, we employed fluorescence recovery after photobleaching (FRAP), a technique that measures protein mobility in live cells. In the FRAP experiments, we used a laser to bleach a small region of mature adherens junction, resulting in the loss of mCherry fluorescence in that area. We then measured the time-dependent recovery in fluorescence signal which occurs due to the exchange of Pcad molecules (Figure 4f). Since previous studies demonstrate that fluorescence recovery in mature adherens junctions primarily arise due to endocytosis and occur on the minute time scale^26,27^, we monitored fluorescence recovery for ∼ 3 min. Fluorescence recovery was fitted to an exponential equation, and the maximum fraction of fluorescence recovery (Figure 4g, ‘mobile fraction’) and the half recovery time (Figure 4g, ‘t_1/2_’) was obtained from the fitted curve (Figure 4g, top panel). The maximum fraction of fluorescence recovery indicates the proportion of mobile proteins in that region with higher recovery signifying increased dynamics at the cell-cell junction while t_1/2_ serves as a proxy for protein turnover rate, with shorter recovery time signifying more rapid turnover. For the A431 cells expressing WT Pcad, the mobile fraction was 38.7 ± 10.2% (Figure 4g, bottom left panel, ‘−CQY’), whereas cells treated with CQY684 Fab exhibited a higher mobile fraction of 52.5 ± 11.4% (Figure 4g, bottom left panel, ‘+CQY’). Consistent with previous results^7^, cells expressing W2A Pcad also had a high mobile fraction (67.7 ± 12.4%; Figure 4g, bottom left panel, ‘W2A’). Similarly, both the W2A mutant and the WT +CQY had a shorter t_1/2_ (35.9 ± 15.2 s and 41.9 ± 13.3s respectively) compared with WT Pcad -CQY (56.0 ± 18.1 s; Figure 4g, bottom right panel). Based on these FRAP results, we concluded that CQY684 increases the dynamics of cell-cell junctions by trapping Pcad in an X-dimer conformation, similar to the W2A mutant. Taken together, these results indicate that X-dimer formation results in the dissociation of p120 which facilitates cadherin endocytosis and weakens Pcad-mediated cell-cell adhesion.

## Discussion

Here, we describe a novel outside-in signaling mechanism that transduces cadherin ectodomain conformation to cytoplasmic signaling events. Specifically, we show that the Mab CQY684 traps and stabilizes the Pcad X-dimer conformation. This results in the dissociation of p120-catenin from the cadherin cytoplasmic region and subsequently triggers Pcad endocytosis. We anticipate that this overlooked outside-in signaling mechanism can be exploited for intracellular drug delivery. Rationally designing antibodies that are engineered to trap and stabilize X-dimers would result in internalization of the antibody-cadherin complex and subsequent lysosomal targeting for intracellular drug release.

We have previously described how Ecad-activating monoclonal antibodies 19A11 and 66E8, which bind to the EC1 or EC2 domains, strengthen Ecad S-dimers by forming interactions with the K14 residue, thereby stabilizing the S-dimer binding interface. The K14E point mutation abolishes the strengthening effect of 19A11 and 66E8, emphasizing the critical role of this residue in S-dimer stabilization^11,12^. Additionally, since the K14 residue is involved in cadherin X-dimer formation, 19A11 and 66E8 trap Ecad in the S-dimer state. By preventing the transition from S-dimer to X-dimer, 19A11 and 66E8 may stabilize cadherin on the cell surface and inhibit turnover, thereby strengthening Ecad-mediated cell adhesion. In contrast, CQY684 binds to the Pcad EC1 domain but does not inhibit the formation of either X-dimers or S-dimers. Using cell-free experiments and computational simulations, we have demonstrated that CQY684 stabilizes the Pcad X-dimer interface, selectively strengthening the X-dimer without affecting the S-dimer, thus slowing the transition from X-dimer to S-dimer conformation. This selective strengthening of the X-dimer may be due to the lack of interactions with the K14 residue. The K14 residue could therefore serve as an important epitope for the further design of mAbs that regulate cadherin interactions.

PCA062, a drug-antibody conjugate that uses CQY684 linked to an anti-cancer drug, was previously found to be targeted to lysosomes for therapeutic drug delivery^16^. Our results demonstrate that CQY684 traps Pcad in an X-dimer conformation and subsequently promotes Pcad internalization. PCA062 had entered ‘first-in-human’ phase I clinical trials for treating Pcad-positive tumors. However, this trial was discontinued due to lack of efficacy and reported cancer progression across all participants^17^. One possible explanation for the failure of PCA062 is the overlooked effect of CQY684 disrupting cell adhesion. Loss of cell adhesion occurs during the early stages of cancer metastasis, and disrupting cell adhesion promotes cancer progression^21,28^. To avoid unexpected changes in cell adhesion, future experiments using CQY684 as a drug delivery platform should target cancer cell lines where Pcad is not the predominant cadherin.

In addition to addressing the overlooked effects of CQY684 on cell adhesion, our study resolves several structural inconsistencies regarding the binding of CQY684 to Pcad. It was originally suggested that when bound to CQY684, Pcad dimerizes in a unique asymmetric *trans* binding conformation that inhibits S-dimer formation^16^. However, the crystal structure (PDB 6ZTR) unambiguously demonstrates that CQY684 binds to the Pcad X-dimer conformation. This is in fact an expected result since the Pcad used in the crystal structure possess an N-terminal extension that is known to preferentially form X-dimers^6^. Consequently, the asymmetric binding conformation that was originally proposed^16^ is likely a misinterpretation of the crystal structure. Furthermore, it was claimed that CQY684 disrupts Pcad *cis*-dimer formation due to steric collisions between the antibody binding interface and the *cis*-dimer interface^16^. However, an alignment of CQY684 on previously resolved Pcad *cis* dimer reveals no such steric interference suggesting that CQY684 does not disrupt *cis*-dimer formation (Supplementary Figure S6).

In addition to its therapeutic potential, CQY684 can also serve as a powerful tool for studying the outside-in regulation of cadherins without introducing mutations in the cadherin ectodomain. We have shown that CQY684 traps Pcad in the X-dimer conformation, leading to the dissociation of p120 from the cytoplasmic region, thereby promoting Pcad endocytosis and colocalization with lysosomal markers. However, while the Pcad W2A mutant also shows decreased p120 association, it does not colocalize with the lysosome (Supplementary Figure S7a). This may be due to low proteolytic processing of the cytoplasmic region of the W2A mutant (Supplementary Figure S7b), which is obligatory for cadherin endo-lysosomal trafficking^29^. Therefore, the W2A mutation not only traps cadherin in the X-dimer conformation but may also silence other signaling pathways, confounding studies on the effects of cadherin X-dimers in cells.

Previous studies suggest that X-dimers promote cadherin turnover on the cell surface by binding more weakly than S-dimers^7^. However, this explanation overlooks the outside-in signaling mechanism revealed by our results. Our data suggests that cadherins regulate cell adhesion, not merely by strengthening ectodomain binding, but also by using different binding conformations to signal changes in the association/dissociation of cytoplasmic proteins. Our results indicate that the signaling function of these adhesive conformations could be more crucial than a naïve interpretation based on differential binding strengths in solution.

Our study indicates that the strength of cell adhesion cannot be solely estimated by the amount of cadherins on the cell surface; associated adaptor proteins like p120 must also be considered. To estimate the amount of cell surface Pcad associated with p120, referred to as “effective Pcad,” we combined results from surface biotinylation and Co-IP assays, assuming p120 only associates with surface-localized Pcad (Supplementary Figure S8). We found that effective Pcad levels in both WT +CQY and W2A conditions were half of those in WT -CQY conditions. This reduction in effective Pcad likely explains the disruptions in cell adhesion caused by CQY684 or the W2A mutations, rather than just changes in surface Pcad levels. Notably, WT +CQY has a similar amount of effective Pcad as W2A but stronger cell adhesion, possibly because WT +CQY forms stronger X-dimers than W2A, and also WT +CQY can still forms S-dimers, whereas W2A cannot.

While our data demonstrates that endocytosis is the primary contributor to Pcad turnover, we cannot eliminate the contribution of Pcad diffusion towards increased junctional dynamics. Previous FRAP measurements on adherens junctions demonstrate that rapid endocytosis is the overwhelming contributor to dynamics in mature Ecad junctions^26^. In contrast, when adherens junctions are disrupted by chelating Ca^2+^, Pcad diffusion increases significantly^26,30^. To decrease the effect of non-physiological diffusive contributions to Pcad dynamics, we performed FRAP experiments on cells that were cultured for at least 24 hours to form mature junctions, and we performed all experiments in the presence of Ca^2+^. Since endocytosis was shown to occur on the minute timescale^26,27^, we monitored fluorescence recovery for ∼ 3 mins to capture rapid endocytosis events.

Our study reinforces the role of p120 as a primary regulator of cadherin dynamics on the cell surface. Unlike previous studies^9,10^ that mutated the cadherin cytoplasmic regions to prevent p120 binding, we used an antibody targeting the ectodomains of Pcad. This approach promoted p120 dissociation by trapping the Pcad ectodomains in an X-dimer conformation. Our results establish a link between cadherin extracellular binding conformations and intracellular signaling. Recent studies suggest that changes in the phosphorylation state of p120 regulates cell adhesion. Specifically, antibodies 19A11 and 66E8, which strengthen Ecad S-dimers and enhance Ecad-mediated cell adhesion, were shown to cause p120 de-phosphorylation in Colo-205 cells^14,15^. It is therefore possible that the phosphorylation states of p120 is also altered due to X-dimer formation, which results in the dissociation of p120 from the Pcad cytoplasmic region. However, dissociation of p120 can also be induced by changes in the phosphorylation state of the cadherin cytoplasmic region^31^, or by the association of non-phosphorylated Numb and/or adaptor protein 2 (AP-2) to a dileucine motif on the cadherin cytoplasmic region^9,32^.

Since both cadherin phosphorylation sites and the cadherin dileucine motif are adjacent to the p120 binding site, we reasoned that X-dimer formation may allosterically induce conformational changes on the Pcad cytoplasmic region that expose these sites. A previous model suggests that since the S-dimer binding interface is at the tip of the outermost EC1 domain, cell-cell junctions are wider when cadherins form S-dimers^7^. In contrast, since the X-dimer binding interface is at the junction of the EC1-EC2 domain, the intercellular gap narrows when cadherins form X-dimers^7^. Consequently, we hypothesized that X-dimer formation generates a tensile stress on the cadherin cytoplasmic region which induces a conformational change that subsequently exposes phosphorylation sites and the dileucine motif (Figure 5a). To test this model, we performed MD and SMD simulations on the crystal structure of the cadherin cytoplasmic juxtamembrane (JMD) core region (residues 758-775) interacting with portion of p120 (residues 359-511)^25^. Although the cadherin phosphorylation sites (residues 755 and 756), and AP-2 binding sites (residues 743 and 744) are not resolved in the crystal structure, they are immediately adjacent to the p120 binding site which suggests that association of cadherin with p120 may sterically hinder access to these sites. We first used 20 ns MD simulations to equilibrate the structures and observed that the cadherin cytoplasmic tail was tightly bound to p120 via an extensive network of hydrogen bonds (∼15 hydrogen bonds, Figure 5b). To mimic a tensile force on the cadherin cytoplasmic tail due to X-dimer formation, we performed SMD simulations where we fixed the C-terminal residue of the cytoplasmic regions and applied a constant stretching force to the N-terminal residue (Figure 5b). Upon application of a tensile force, the hydrogen bond network between p120 and the cadherin cytoplasmic region decreased over time (Figure 5c), suggesting that the stress on the cytoplasmic region weakens these interactions, thereby exposing both the phosphorylation sites and dileucine motif. Unfortunately, since the p120 phosphorylation sites (residues 252, 268, 288, 310, and 312) are unsolved in the crystal structure, the effect of X-dimer formation on p120 phosphorylation remains unclear. It is also important to note that although the model we propose is supported by simulations, it remains to be experimentally validated.

**Figure 5.**
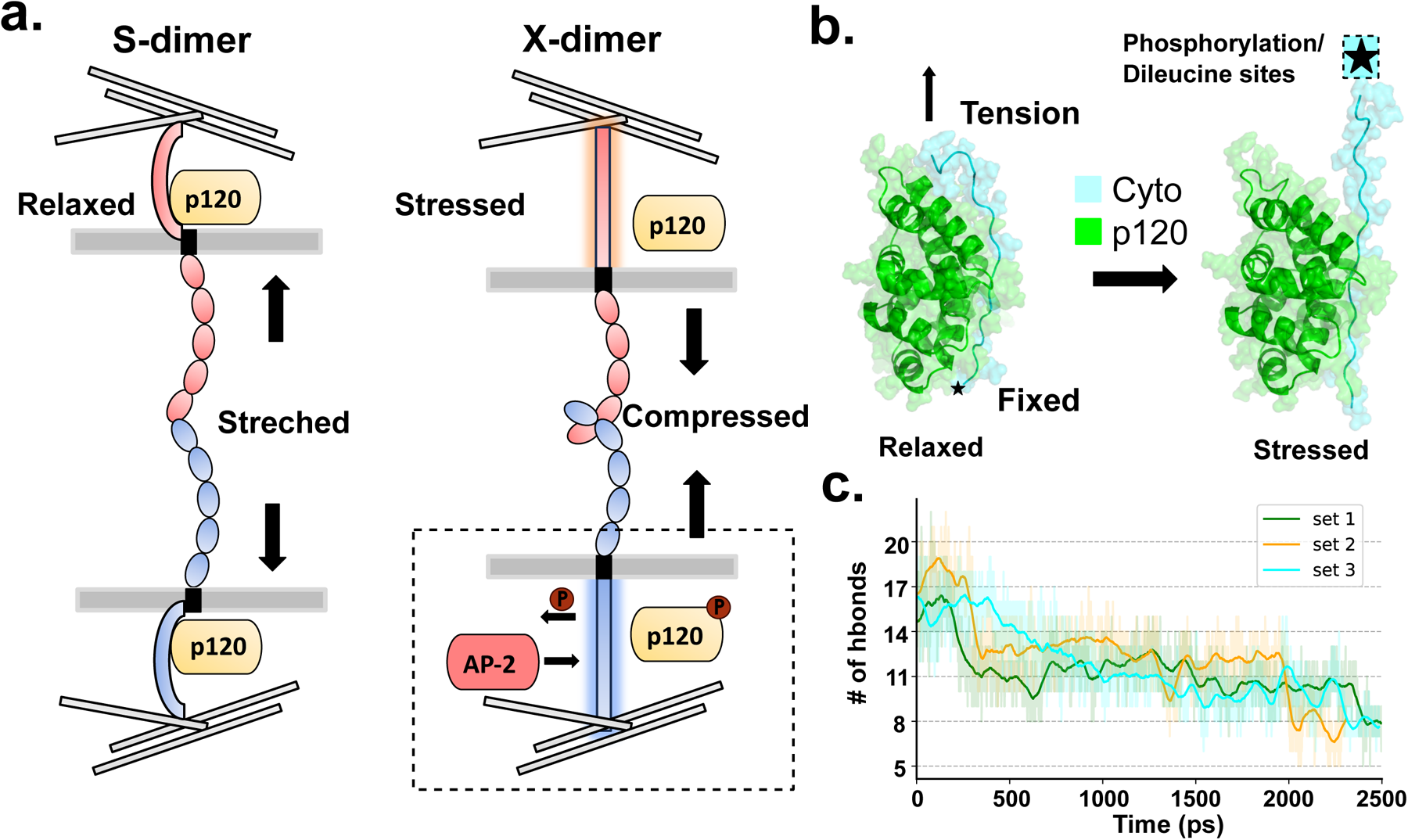
Proposed model for X-dimer induced p120-catenin dissociation. (a) Schematic of proposed model. Formation of X-dimer compresses the cell membrane, which subsequently loads the cytoplasmic region and changes its conformation. This triggers p120 dissociation by either changing the phosphorylation state of the p120/cadherin, or by the association of AP-2. (b) (c) SMD simulations show that loading the cadherin cytoplasmic region changes its conformation, which weakens p120 binding and exposes both cadherin phosphorylation sites and the AP-2 binding site. (b) Structures observed during the SMD. The cadherin cytoplasmic region is in cyan while p120 is in green. (c) The number of hydrogen bonds formed between p120 and the cadherin cytoplasmic region decreases as the cadherin cytoplasmic region is loaded.

In summary, our study shows that CQY684 traps Pcad in an X-dimer conformation and stabilizes this adhesive structure. Formation of stable X-dimers leads to the dissociation of p120 from the cadherin cytoplasmic region, thereby promoting Pcad turnover and targeting Pcad to the lysosome. Our results suggest an outside-in signaling mechanism that can be exploited by anti-cadherin antibodies for intracellular drug delivery.

## Materials and Methods

### Purification of human WT, W2A, and K14E Pcad ectodomains and CQY684 Fab

The ectodomain of Pcad (1-654) was cloned with a C-terminal Avi-tag and 6XHis-tag and incorporated into pcDNA3.1(+) as previously described^11,12^. The mutant W2A and K14E Pcad were constructed using the Q5 site-directed mutagenesis kit (New England Biolab). The plasmids were transfected into Expi293 suspension cells with the ExpiFectamine transfection kit (ThermoFisher Scientific). After 5 days of expression, the conditioned media which contain Pcad ectodomains were collected, and a protease inhibitor tablet (ThermoFisher Scientific) was added to the media. The WT and mutant Pcads were then affinity purified with Ni-NTA agarose beads (Qiagen) in a gravitational column. The Ni-NTA beads were first washed with binding buffer (pH 7.5, 20 mM HEPES, 500 mM NaCl, 1 mM CaCl_2_), followed by washing with biotinylation buffer (pH 7.5, 25 mM HEPES, 5 mM NaCl, 1 mM CaCl_2_). The Ni-NTA-bound Pcads were then biotinylated using the BirA enzyme (BirA 500 kit, Avidity). The biotinylated proteins were eluted with elution buffer (pH 7.5, 500 mM NaCl, 1 mM CaCl_2_, 200 mM imidazole). The eluted proteins were dialyzed to remove imidazole using storage buffer (pH 7.5, 10 mM Tris-HCl, 100 mM NaCl, 10 mM KCl, 2.5 mM CaCl_2_). 20% Glycerol was added into the protein sample before the samples were flash-frozen in liquid nitrogen and stored at -80°C.

The sequence of CQY684 Fab was previously described^16^ and deposited in PDB (access code 6ZTR). CQY Fabs were recombinantly generated by cloning the variable regions of the heavy and light chains into the human IgG Fab backbone by GenScript.

### Single-molecule AFM experiments

The single-molecule AFM experiments were performed using the previously described protocol^11,12,18^. Briefly, the cantilever (Hydra-2R-50N, AppNano) and glass coverslips (CS) were submerged in Piranha solution (25% H_2_O_2_, 75% H_2_SO_4_) overnight and washed with DI water twice. The CS was also treated with 1 mM KOH and washed with DI water twice. The cantilever and CS were then washed with acetone and silanized with 2% 3-aminopropyltriethoxysilane (Millipore Sigma) in acetone for 30 minutes at room temperature. After that, the cantilever and CS were functionalized with 10% biotin-PEG-Succinimidyl Valerate (MW 5000, Laysan) and 90% mPEG-Succinimidyl Valerate (MW 5000, Laysan) in 100 mM NaHCO_3_ and 600 mM K_2_SO_4_ for at least 4 hours of incubation and followed by washing with DI water.

Prior to the AFM experiments, the cantilever and CS were first blocked with 10 mg/mL BSA for 1 hour at room temperature and then incubated with 0.1 mg/mL streptavidin (Invitrogen) for 30 minutes at room temperature. Finally, the functionalized cantilever and CS were incubated with 200 nM of purified proteins for 90 minutes.

All AFM experiments were carried out using an Agilent 5500 AFM system with a closed-loop scanner. The single-molecule force measurements were performed in a Tris buffer (pH 7.5, 10 mM Tris-HCl, 100 mM NaCl, 10 mM KCl, 2.5 mM CaCl_2_). The spring constants were calculated using the thermal fluctuation method. To filter out the unspecific binding events, all recorded PEG stretching force curves were fitted into a worm-like-chain (WLC) model using the least-squares fitting method. The populations of unbinding forces were fitted using a Gaussian mixture model through binning based on the Freedman-Diaconis rule. The Bayesian Information Criterion (BIC) was used to determine the optimal number of Gaussians in the unbinding force distributions.

### Bead aggregation assay

Biotinylated ectodomains of WT, W2A, and K14E P-cad at a concentration of 200 nM were mixed with Dynabeads™ MyOne™ Streptavidin C1 (Thermo Fisher Scientific) and incubated for 30 minutes in binding buffer (10 mM Tris-HCl, 150 mM NaCl, 0.05% Tween 20, 1 mg/ml BSA, pH 7.5) at 4°C with gentle shaking at 90 RPM. Following incubation, the Dynabeads were washed and then transferred to either Calcium binding buffer (10 mM Tris-HCl, 150 mM NaCl, 5 mM CaCl_2_, 0.05% Tween 20, 1 mg/ml BSA, pH 7.5) or CQY684 binding buffer (100nM CQY684 Fab, 10 mM Tris-HCl, 150 mM NaCl, 5 mM CaCl_2_, 0.05% Tween 20, 1 mg/ml BSA, pH 7.5) and vigorously rotated at room temperature for 2 hours. After incubation, 10 µl of the sample was placed onto a sample slide and imaged using a 10X objective with an Olympus CX40 microscopy system. Analysis was performed using imageJ2 with a 30-pixel cutoff.

### Molecular Dynamics (MD) and Steered Molecular Dynamics (SMD) simulations

MD simulations were conducted on the FARM high-performance computing cluster at the University of California, Davis, using GROMACS 2022.3 as previously described^11,12^. The simulations utilized the OPLS-AA/L force field and the TIP4P water model with a 10 Å radius cut-off for Van der Waals and electrostatic interactions. Electrostatic energy was calculated using the particle mesh Ewald method with a 0.16 nm grid spacing. The Pcad X-dimer structures with EC1 and EC2 domains, and CQY bound X-dimer complex structures were prepared by generating the symmetry mates in PyMOL using the crystal structure PDB: 6ZTR. The structures were then prepared for simulations using PDBFixer.

At the start of the MD simulations, the Pcad X-dimer structures either bound or not bound to CQY684 were placed in the center of a dodecahedral box, ensuring every atom was at least 1 nm away from the boundary. The system was relaxed with energy minimization and stabilized with equilibration under isothermal-isochoric and isothermal-isobaric conditions using a modified Berendsen thermostat and Berendsen barostat. The box was filled with water molecules and neutralized with charged ions (100 mM NaCl, 4 mM KCl, and 2 mM CaCl_2_).

After stabilization, a 60 ns MD simulation was performed with 2-fs integration steps, with the protein structure typically reaching equilibration after approximately 20 ns (Supplementary Figure S4). Using the gmx rmsf module, the C-α RMSF of residues 1 to 100 in the EC1 domain was monitored from the backbone RMSD of the structure relative to the initial frame. To calculate the RMSF of the structure during the MD simulations, we used gmx rmsf module.

Simulations for p120 bound to the Ecad cytoplasmic region was constructed from the crystal structure PDB: 3L6X. The protein complex was similarly prepared as described above. However, only 20ns MD simulations were performed.

The constant force SMD simulation were performed on the FARM high-performance computing cluster. Using the final frame from the MD simulation, we obtained an interacting Pcad protein structure and placed it in the center of a rectangular box, aligned with the box’s longest axis, ensuring no atom was closer than 1 nm to the boundary (dimensions: 30 × 12 × 8 nm for the - CQY conditions; 30 × 15 × 15 nm for the +CQY condition). The system, containing approximately 38,000 atoms for the -CQY condition and approximately 88,000 atoms for the +CQY condition, was relaxed and equilibrated using the same method as in the MD simulation, but without the isothermal-isochoric step.

At the start of the SMD simulation, one C-terminus of the interacting Pcad protein pair was fixed while the other C-terminus at residue 213 was pulled along the longest axis of the box with a constant force of approximately 665 pN (400 kJ⋅mol−1⋅nm−1). To evaluate the stability of the structure throughout the SMD simulation, the gmx sasa module was used to calculate the ΔSASA (ΔSASA = SASA [protein A] + SASA [protein B] − SASA [protein A + protein B]), which estimated the change in the Pcad interfacial binding area.

### Generation of stable cell lines

WT-Pcad and W2A-Pcad full-length plasmids fused with mCherry at the C-terminus were introduced into A431 Ecad/Pcad double knockout cell line. Using the limiting dilution-culture method, a single clone of each rescue cell line was acquired. The monoclonal cell line was then expanded and re-selected using fluorescence-activated cell sorting (FACS). A single cell was again seeded onto collagen-treated 96-well plates, completing the generation of stable monoclonal A431 Pcad-mCherry rescued cells. The rescued cells were cultured in Gibco high glucose Dulbecco’s Modified Eagle Medium (DMEM) containing 10% fetal bovine serum (FBS) and 1% penicillin-streptomycin.

### Cell Aggregation Assay

A431 Pcad-mCherry cells were detached using 0.05% trypsin diluted with calcium and magnesium-free solution (CMFS). To prevent the digestion of Pcad on the surface, 1 mM CaCl_2_ was added to the trypsin buffer. After 30 minutes of trypsinization, the cells were suspended, and the reaction was stopped by adding suspension buffer (Life Science Ca-free DMEM with 10% Ca-free FBS) to recollect all suspended cells. The cells were washed again with suspension buffer and diluted to a concentration of 1 × 10^5^ cells/ml. Then, 500 µL of suspended cells were added to each well of a 24-well plate with the supplementation of 2 mM CaCl_2_ or 4 mM EGTA in the presence or absence of CQY684. The 24-well plates were pretreated with 0.5% BSA overnight at 4 °C to prevent cell attachment to the bottom of the plate and were washed three times with PBS prior to start of the experiment. Cells were incubated at 37 °C and 100 rpm for 2 hours. Brightfield images were acquired with an Olympus CKX41 imaging system. Analysis was performed using imageJ2. Images were first converted to 8 bit format, and pixels corresponding to single cells were picked (single cells correspond to threshold of 40 pixels). Only aggregates that contain more than 5 cells were used in the analysis.

### Monolayer dispersion assay

The cells were seeded in 12-well plates in the presence or absence of CQY Fab at a concentration of 1 × 10^5^ cells/ml. After 48 hours of incubation, the cell monolayer was washed twice with high-glucose DMEM to remove any floating cells and then incubated with 1.2 U/ml Dispase II (Millipore Sigma) in PBS with 1 mM CaCl_2_ for 1 hour at 37 °C [6]. After detachment, the lifted cell monolayer was placed on an orbital shaker and subjected to a shaking force of 200 rpm for 40 minutes. The fragmentation of the cell monolayer was imaged and analyzed using a Leica confocal imaging system.

### Immunofluorescence confocal imaging

Cells were plated on a Bovine Collagen I (R&D Systems) coated glass coverslip and cultured in the presence or absence of CQY684 Fab until reaching 70% confluence. The cultured cells were fixed with an ice-cold 1:1 acetone/methanol mixture for 10 minutes at 4 °C. After fixation, the cells were washed three times with PBS and blocked with 5% BSA buffer. The blocked samples were then incubated with a 1:100 dilution of primary antibody for 1 hour, followed by a 1:500 dilution of secondary antibody for 30 minutes at room temperature. Images used to quantify internalization were captured with a Leica confocal microscopy system. Colocalization analysis for Pearson’s coefficients was carried out using ImageJ2 module colo2 with Costes threshold regression method; PSF set to 3 and Costes randomization set to 50.

### Fluorescence Recovery After Photobleaching (FRAP)

To prepare samples for FRAP experiments, cells were seeded onto Cellvis 35 mm dishes coated with Cell-Tak (Fisher Scientific) and cultured for at least 24 hours. During incubation, 200nM CQY684 was added to the A431 cells expressing WT Pcad for the ‘+CQY’ conditions. Subsequently, cells were transferred to a Leica Stellaris 5 confocal microscopy system equipped with an FRAP module. Imaging was performed using a 63X oil immersion objective with a 1 Airy unit pinhole. mCherry fused to Pcad was excited using a 568 nm argon laser, with 2% laser intensity for pre- and post-bleaching imaging, and 30% laser intensity during bleaching. The FRAP protocol consisted of acquiring 10 pre-bleach frames, followed by 10 frames during bleaching, and 200 post-bleach frames (863 ms exposure per frame). Mean integrated fluorescence intensity within each region of interest (ROI) was calculated and normalized to the pre-bleaching value. Background measurements from the same ROI were subtracted during calculations. The recovery phase of the fluorescence signal was fitted to a single exponential equation: 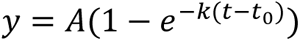. Only R^2^ values above 0.7 were used in subsequent analysis.

### Co-Immunoprecipitation

To prepare the cell lysate for co-immunoprecipitation (Co-IP), cells were seeded in a T75 tissue culture flask (Thermo Fisher Scientific) in the presence or absence of CQY684 Fab. After 48 hours of incubation, the cells were washed three times with PBS and scraped off with 1 ml of PBS. The cells were then resuspended in M2 lysis buffer (pH 7.5, 50 mM Tris-HCl, 150 mM NaCl, 1% SDS, 1% Triton-X 100) with 1% protease inhibitor cocktail (Sigma-Aldrich) and 0.1% Benzonase (Millipore Sigma). The resuspended cell solution was flash-frozen in liquid nitrogen and incubated on ice for 30 minutes. The cell lysates were then sonicated for two 1-minute cycles to ensure full release of proteins. After the lysis process, the cell lysates were centrifuged at 13,000 rpm for 10 minutes at 4 °C to remove cell debris, and the protein concentration was determined using the Bio-Rad DC protein assay with BSA as the standard.

Prior to the co-IP experiments, magnetic protein G beads (Thermo Fisher Scientific) were incubated with anti-mCherry rabbit mAb #43590 (Cell Signaling Technology) at room temperature for 10 minutes. Then, 500 µl of cell lysate was added to the antibody-bound beads and incubated for 30 minutes with rotation at room temperature. The beads were washed three times with washing buffer (Thermo Fisher Scientific). The antigen proteins were eluted by incubating the beads with elution buffer (Thermo Fisher Scientific) for 2 minutes with rotation at room temperature.

### Surface biotinylation assays

The surface biotinylation was performed as previously described^31,33^. In brief, a confluent monolayer of cells formed in the presence or absence of CQY Fab was washed three times with cold PBS. Cell surface proteins were incubated for 10 minutes at 4 °C with EZ-link Sulfo-NHS-SS-Biotin (Thermo Fisher Scientific). Then, 500 µL of quench buffer (Thermo Fisher Scientific) was added to stop the reaction, and the cells were recollected. The cells were lysed with M2 lysis buffer, and the lysates were incubated with NeutrAvidin agarose (Thermo Fisher Scientific) for 60 minutes at room temperature with end-over-end mixing using a rotator. The agarose beads were then washed three times with washing buffer (Thermo Fisher Scientific) and eluted by incubating the beads with SDS-PAGE sample buffer containing 50 mM DTT. The amount of protein in the cell lysate and elution was measured from western blots.

### Western Blots

Purified human WT and mutant W2A/K14E Pcad ectodomains, cell lysates, and eluted proteins from co-IP and surface biotinylation experiments were prepared by boiling at 95 °C for 5 minutes in SDS-PAGE sample buffer (Bio-Rad, 90% 4X Laemmli Sample Buffer + 10% 2-Mercaptoethanol). The denatured samples were run on 4–15% polyacrylamide gels (Mini-PROTEAN TGX Precast Protein Gels, Bio-Rad) at 200V for 30 minutes and then transferred to a PVDF membrane (Bio-Rad) at 200 mA for 1 hour on ice. The membrane was then blocked with 5% blocking buffer (PBS + 0.1% Tween 20 + 5% blotting-grade blocker) for 1 hour and washed three times with PBST. After a 1-hour incubation with a 1:1000 dilution of the primary antibody and a 1:5000 dilution of the secondary antibody, the protein was detected using WesternBright ECL HRP substrate (Advansta).

## Acknowledgements

This research was supported by the National Institute of General Medical Sciences of the National Institutes of Health (R01GM133880). We thank Prof. Sergey Troyanovsky (Northwestern University) for sharing the A431 Ecad/Pcad double knockout cells. FACS was performed at the Flow Cytometry Shared Resource funded by the UC Davis Comprehensive Cancer Center Support Grant (CCSG) awarded by the National Cancer Institute (NCI P30CA093373).

## Author Contributions

B.X., S.X., and S.S. designed the research. B.X., and S.X. performed the research and analyzed the data. S.S. directed the research. B.X., S.X. and S.S. wrote the paper.

