## Supplementary Information for "Outside-in engineering of cadherin endocytosis using a conformation strengthening antibody"

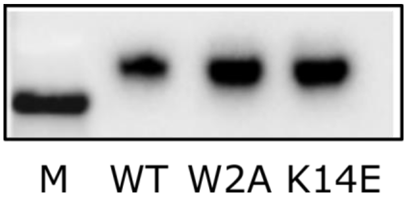


**Supplementary Figure S1.** Western blots confirm that CQY684 recognizes Wild type (WT), W2A, and K14E Pcad. The molecular weight marker (M) corresponds to a molecular weight of 75 kD.

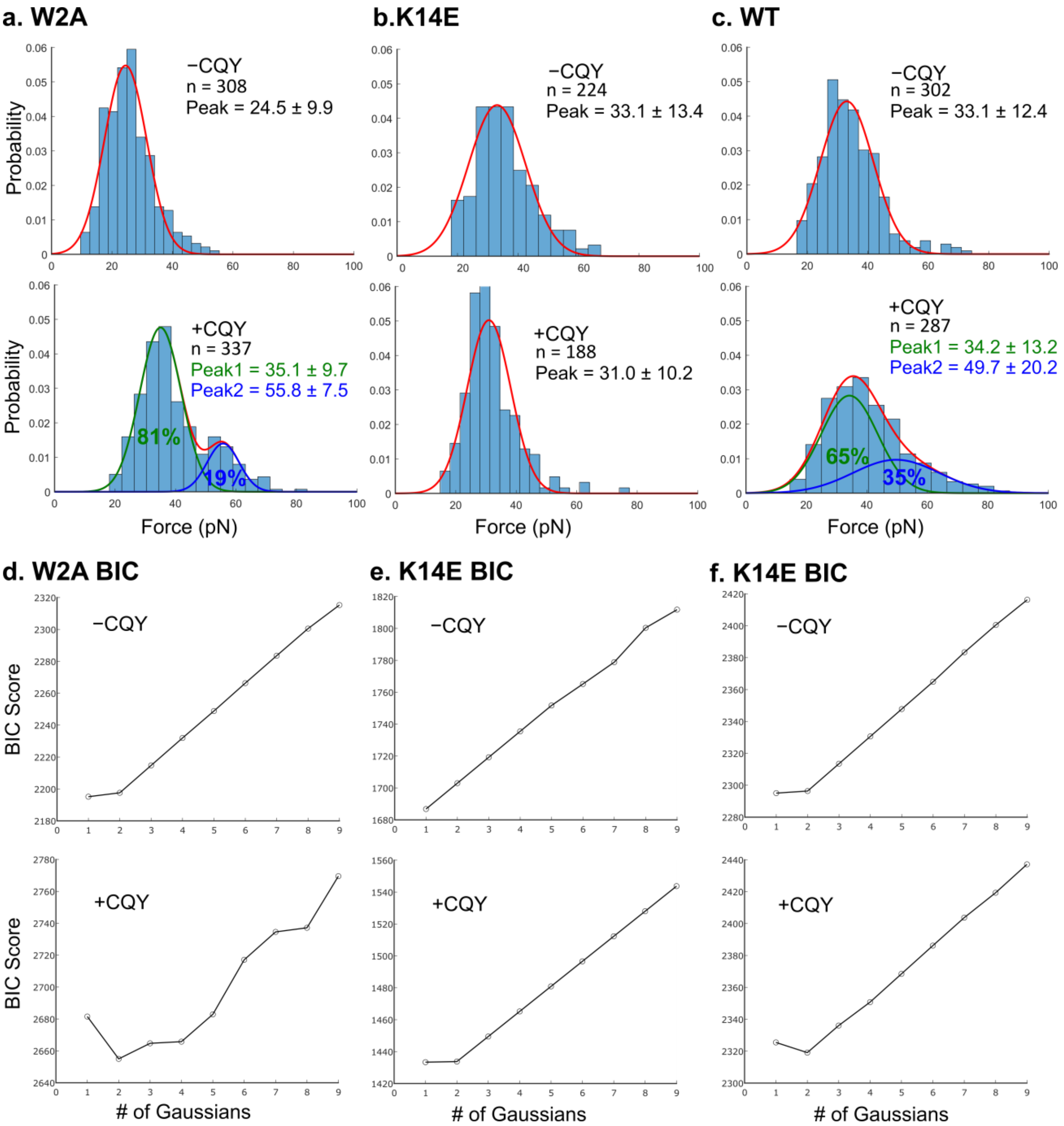


**Supplementary Figure S2. AFM histograms and the corresponding Bayesian Information Criterion (BIC) test.** Histograms of unbinding forces with or without CQY684 Fab for the experiments using (a) W2A, (b) K14E, and (c) WT Pcad. (d) (e) (f) Corresponding BIC testing shows that all force distributions are best described by a single Gaussian distribution expect the W2A + CQY and WT + CQY conditions, which are best described by two Gaussian distributions.

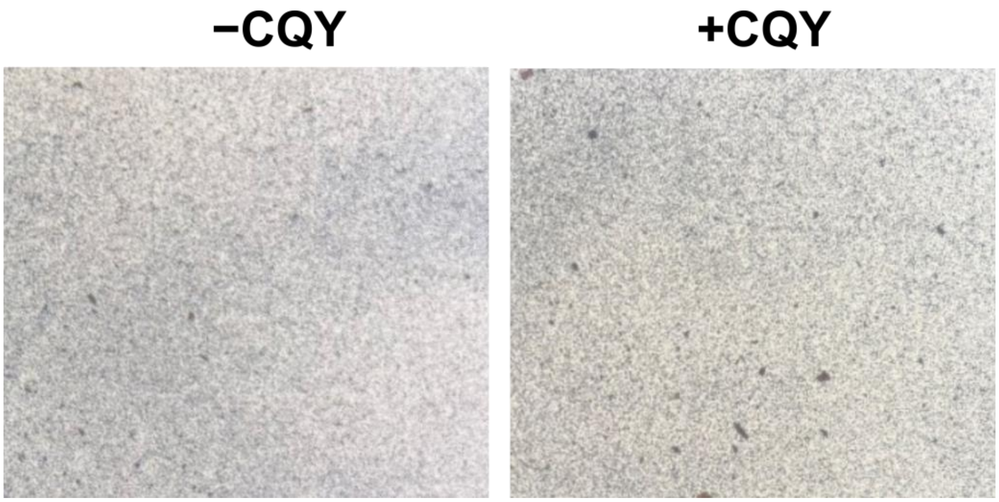


**Supplementary Figure S3. Addition of CQY684 does not alter K14E Pcad bead aggregation.** Significant bead aggregation was not observed for K14E Pcad with or without CQY684.

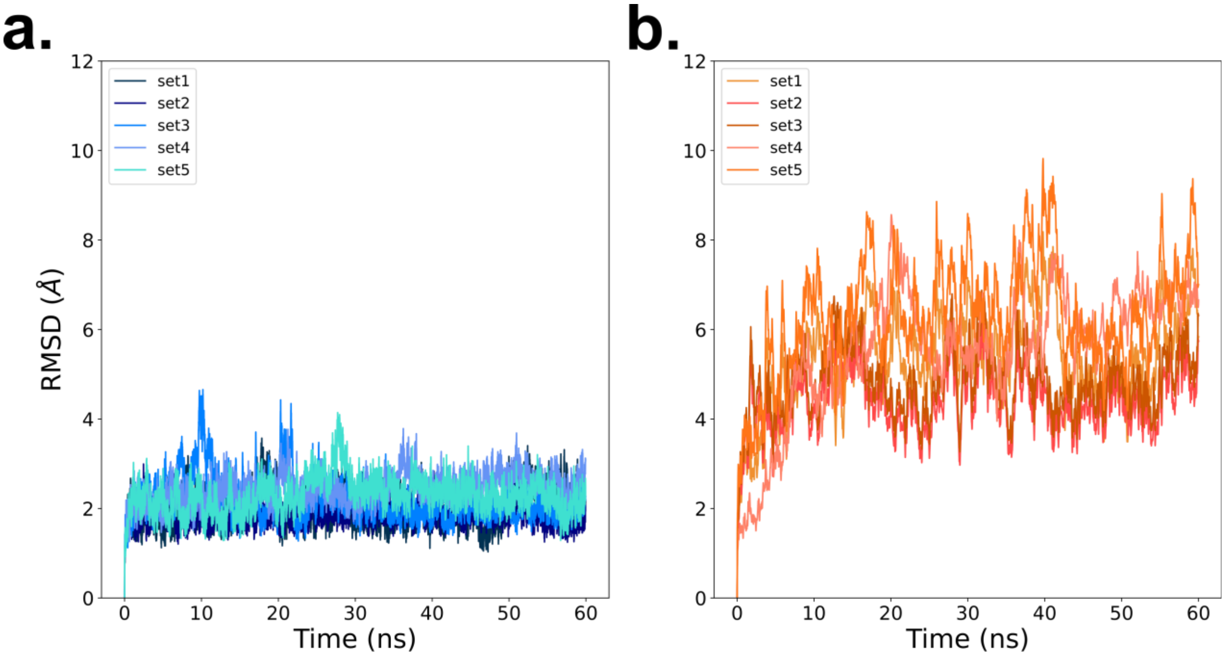


**Supplementary Figure S4. Protein backbone RMSD in MD simulations relative to the initial structures at the start of simulation.** RMSD values measured for each set in the two conditions (a) −CQY condition, and (b) +CQY condition. Stabilization of RMSD values suggest that structures are well equilibrated.

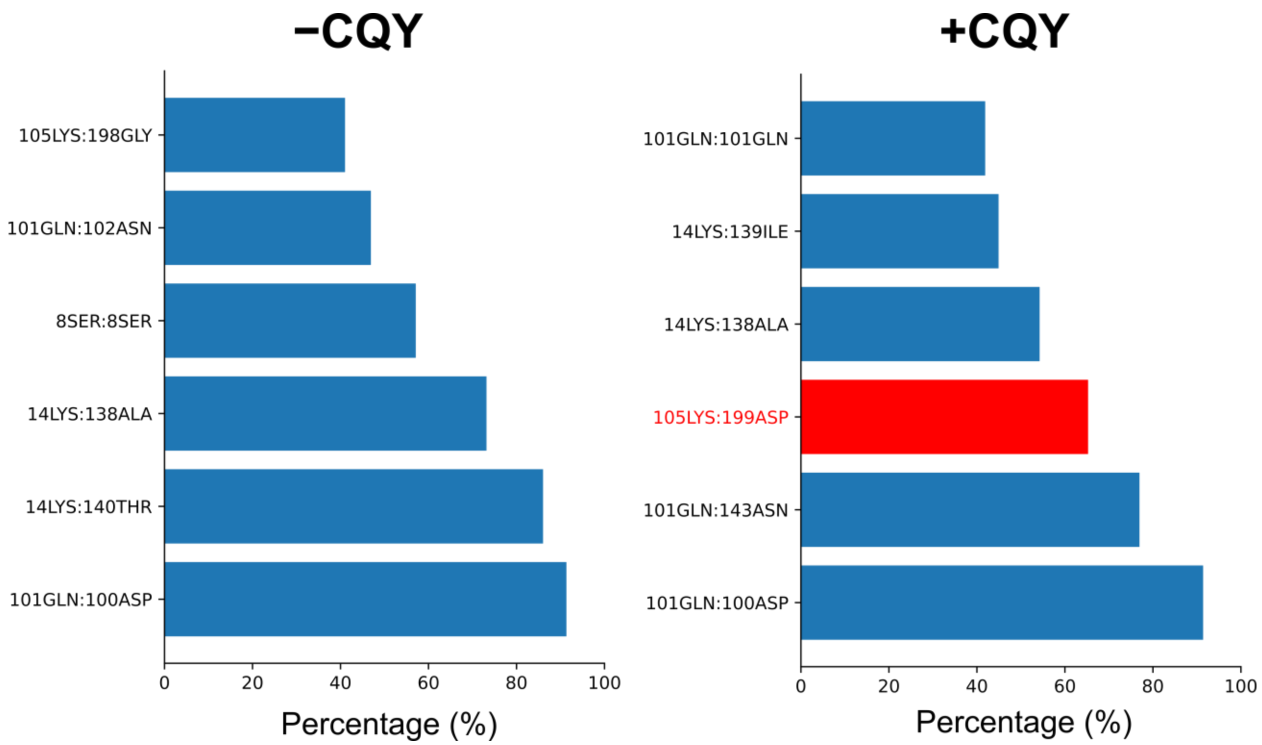


**Supplementary Fig. S5. Electrostatic interactions between Pcads in an X-dimer change upon interaction with CQY684.** Bar plots of all the electrostatic interactions which include salt bridges and hydrogen bonds that are persistent over 40% of the total simulation time are shown above. The addition of CQY684 introduced a novel salt bridge 105LYS:199ASP, which is highlighted in red.

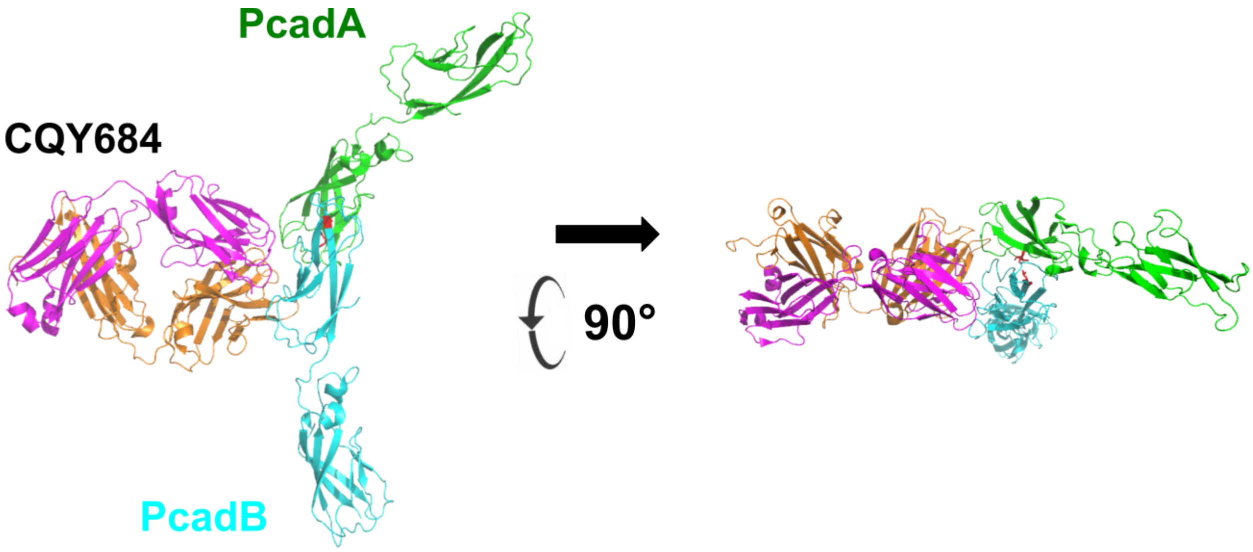


**Supplementary Figure S6. CQY684 binding does not interfere with Pcad *cis* dimer interface.** CQY684 was aligned on a Pcad *cis* dimer (colored in green and cyan, PDB code: 4ZMX). The hydrophobic core of the Pcad *cis* interface, I175 and V81 is highlighted in red. There is no interference observed between CQY684 with Pcad *cis* dimer.

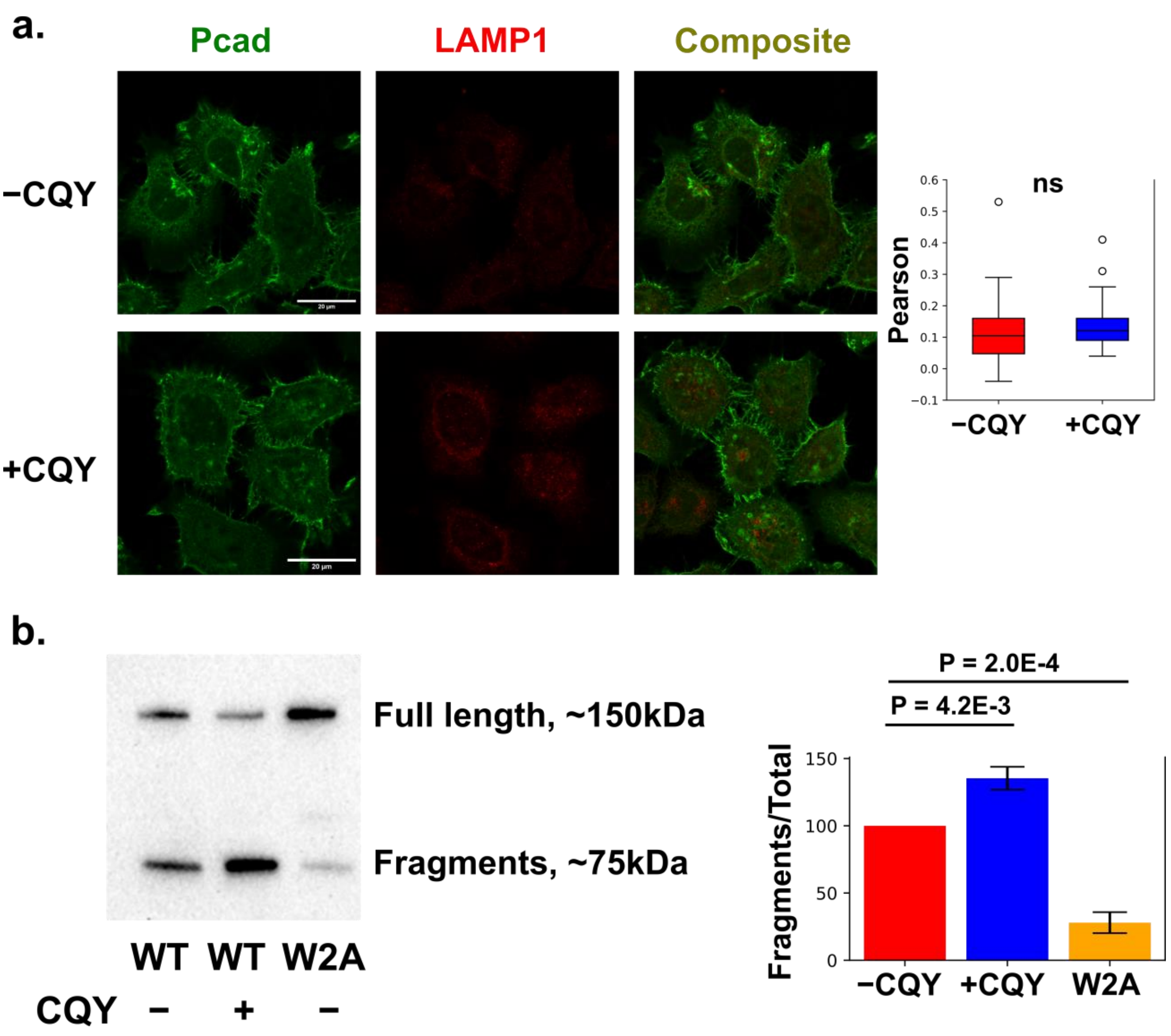


**Supplementary Figure S7. CQY684 does not induce W2A Pcad colocalization with LAMP1 due to low proteolysis activity in Pcad W2A cell line.** (a) Immunofluorescence confocal imaging shows that W2A-Pcad do not colocalize with the lysosomal marker LAMP1 in the presence or absence of CQY684. N = 50 cells for both conditions. Student t-test was performed, and no significant difference observed for colocalization Pearson's coefficients. (b) Left panel: western blots detecting Pcad observed two bands in the cell lysate across the three conditions: 'WT -CQY', 'WT +CQY', and 'W2A'. The top band, which has molecular weight ~150kDa, corresponds to the full length Pcad, while the bottom band, which has molecular weight ~75kDa, corresponds to the proteolytic fragments of Pcad. Right panel: Bar-plot of three replicates. All values are compared to the −CQY conditions. The proteolytic fragments of Pcad increase in the +CQY conditions, corresponding to more endo-lysosomal activity induced by the addition of CQY684, but dramatically decrease in the W2A mutation. This suggests that the W2A mutation silences proteolytic activity, which is required for endo-lysosomal trafficking of Pcad.

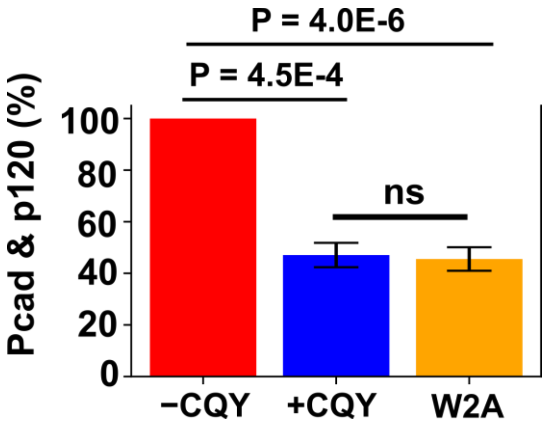


**Supplementary Figure S8. The amount of ‘effective Pcad’ is reduced in both +CQY and W2A conditions.** Bar-plot of three replicates. All values are compared to the −CQY conditions.

**Supplementary Movie.**

Example constant-force SMD simulations. One constant-force SMD example from ‘−CQY’ and ‘+CQY’ conditions. Pcad X-dimers in the ‘−CQY’ condition break significantly faster compared with the +CQY condition. Color scheme: Pcad (blue and red), CQY684 heavy chain (magenta), and CQY684 light chain (orange).
